# scParadise: Tunable highly accurate multi-task cell type annotation and surface protein abundance prediction

**DOI:** 10.1101/2024.09.23.614509

**Authors:** Elizaveta Chechekhina, Vsevolod Tkachuk, Vadim Chechekhin

**Author notes:** Corresponding author Vadim Chechekhin, Elizaveta Chechekhina.

## Abstract

scRNA-seq is revolutionizing biomedical research by revealing tissue architecture, cellular composition, and functional interactions. However, accurate cell type annotation remains a challenge, particularly for rare cell types, with existing automated methods often falling short. Multimodal data, combining mRNA expression and protein markers, improves deep cellular analysis and make functional characterization of complex tissues more accurate. However, it is costly and complex to obtain. We present **scParadise**, a cutting-edge Python framework featuring three tools: **scAdam** for multi-level cell annotation, **scEve** for surface protein prediction, and **scNoah** for benchmarking. scAdam surpasses current methods in annotating rare cell types and ensures consistent results across diverse datasets, while scEve enhances clustering and cell type separation. With scNoah’s advanced metrics, scParadise offers a powerful, fast, and reliable solution for single-cell analysis, setting a new standard in scRNA-seq data processing.

## Introduction

Modern biomedical research increasingly relies on the scRNA-seq method. This method allows us to study the architecture of tissue, its cellular composition, intercellular interactions, and the functional interactions of cells^1^. The crucial step in scRNA-seq analysis is cell type annotation. Manual annotation is a time-consuming process that requires a high level of knowledge about the cellular composition of the tissue under study. This has led to the rapid development of automatic cell type annotation methods^2^. However, the predictive power of existing automatic annotation methods often does not exceed 90%. At the same time, rare cell types (less than 1% of the population) are poorly annotated by most automatic annotation methods.

In addition to mRNA expression, scRNA-seq methods also allow for the abundance of protein markers located on the cell surface. Such multimodal data help solve the problem of clusterization and cell type annotation^3^. Moreover, this approach achieves more accurate quality control and description of the functional state of cells. However, experiments involving multimodal data collection are very expensive and labor-intensive. Consequently, methods for predicting modalities are of significant interest to researchers.

The development of automatic cell annotation methods has led to an unsystematic approach by scientists in validating the results of model performance. Additionally, model testing is often carried out on a single test dataset, which does not reflect the full variability of scRNA-seq data^4,5^. In this regard, there is an increasing need to unify approaches to testing new methods and comparing them with existing ones. In most cases, assessing the quality of a method using one or two metrics (most often accuracy and F1-score) is not sufficient to fully reflect the quality of the model due to the specificity of scRNA-seq data. This requires a tool that allows for a more detailed quality assessment at the level of individual cell subpopulations. In addition, the development of modality prediction methods also requires unification in quality control.

We introduce scParadise, a new Python framework for fast, tunable, high-performance automatic cell type annotation and modality prediction. scParadise includes three sets of tools: scAdam - multi-task multi-class cell type annotation; scEve - modality prediction; scNoah - benchmarking cell type annotation and modality prediction. Additionally, we present pipelines for training and using scParadise models in Python and R.

## Results

### Multi-class cell type prediction using scAdam

To demonstrate the performance of scAdam models, we selected the publicly available CITE-seq PBMC 3’ dataset (Fig. 1A)^6^. scAdam included 3 steps for model training: feature selection, automated dataset balancing, and training (Figure 1A). At the first step, features were manually selected based on the highly variable and marker genes for the cell types. Genes whose expression was uniformly distributed between different cell subpopulations or was detected in single cells of a certain cell type were excluded from the list of trainable features (Suppl. Fig. 1A). We also added marker genes of cell subpopulations not from the list of highly variable genes (Suppl. Fig. 1B). This approach allowed us to select the most significant genes for cell annotation for training the model. Single-cell RNA-seq data were imbalanced in the number of different cell types. In this regard, in the second step, we balanced cell types using the most detailed level of annotation. We undersampled the majority (most represented) cell types (CD14 Mono, NK, CD4 T Naive, etc.) to the average cell count (number of cells divided by number of cell types) using random cell selection. We also oversampled the rare (least represented) cell types (ASDC, ILC, HSPC, etc.) by picking cells at random. This approach allowed us to increase balanced accuracy (+4.5%), F1-score (+1.9%), geometric mean (+2.7%), and index balanced accuracy of the geometric mean (+4.6%) with a slight (−0.5%) decrease in accuracy (Fig. 1C-D, Suppl. Fig. 1C) in the case of celltype_l2 classification. This also allowed us to increase some metrics (sensitivity, precision, F1-score) for several cell types such as ILC, double-negative T cells, ASDC, and B naive lambda cells (Suppl. Fig. 4-5).

**Figure 1.**
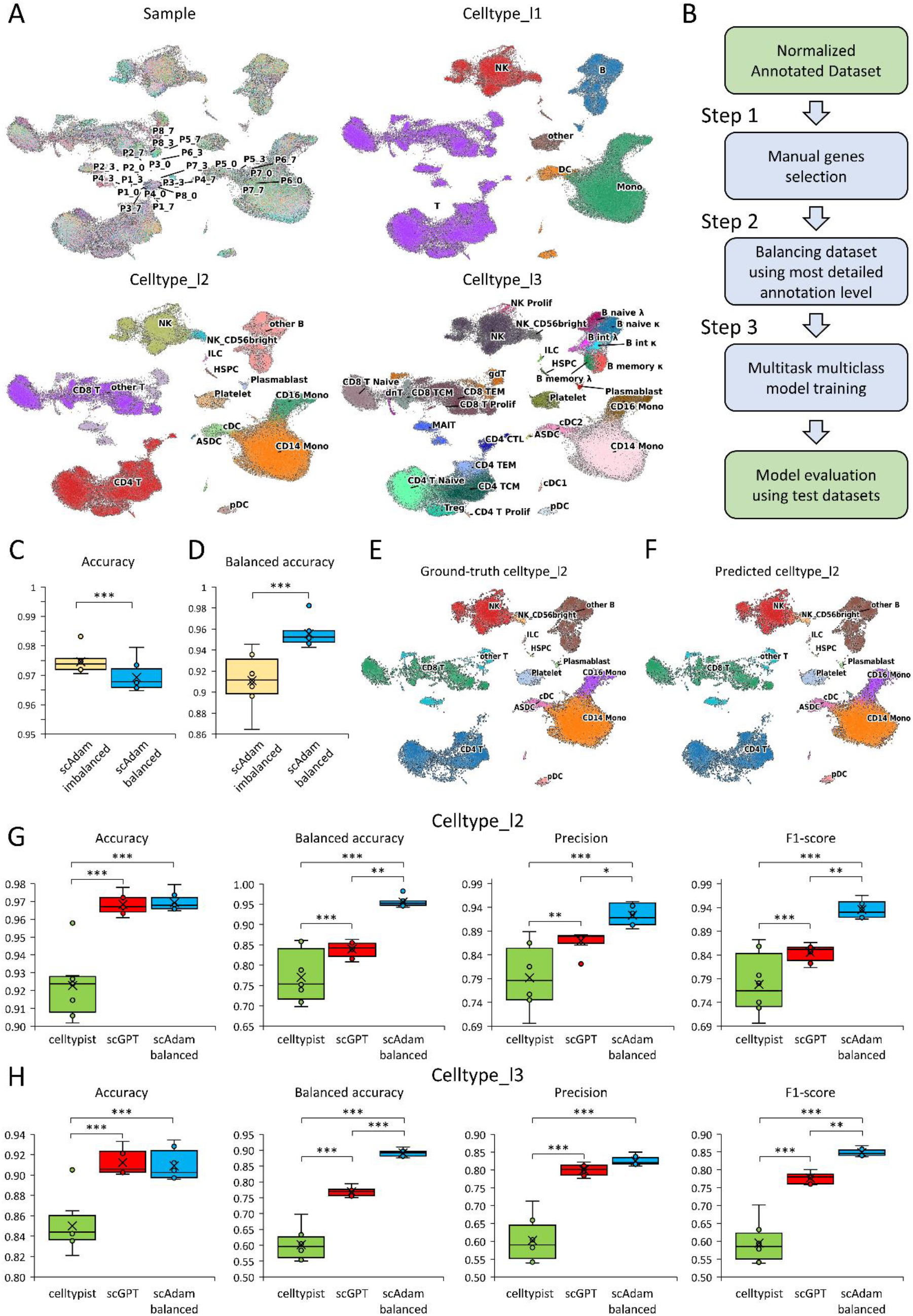
scAdam model training and comparison with other AI-based annotation methods. A. WNN- integrated CITE-seq PBMC 3’ dataset annotated on 3 levels (celltype_l1 – least detailed, celltype_l3 – most detailed). B. Pipeline of scAdam model training and evaluation. C-D. Box plots of accuracy (C) and balanced accuracy (D) between imbalanced and balanced datasets. Paired T-test. E. UMAP visualization of cells colored by ground-truth cell types (celltype_l2) in the PBMC test dataset. F. UMAP visualization of cells colored by scAdam predicted cell types (predicted celltype_l2). G-H. Box plots comparison of celltypist, scGPT and scAdam models in average accuracy, balanced accuracy, precision and f1-score using 8 test datasets (G – celltype_l2, H – celltype_l3). One Way RM ANOVA with post-hoc Tukey test. Median (line), mean (cross), n = 8 (dots), * p<0.05, ** p<0.01, *** p<0.001.

For training the model, we selected 8 datasets out of 24. The remaining datasets were used as test data. Figure 1 show the ground truth (E) versus predicted (F) cell types (celltype_l2) by the scAdam model. Other annotation levels (celltype_l1, celltype_l3) with predicted probabilities (higher values meaning higher probabilities of accurate annotation) were presented in Suppl. Fig. 2A.

**Figure 2.**
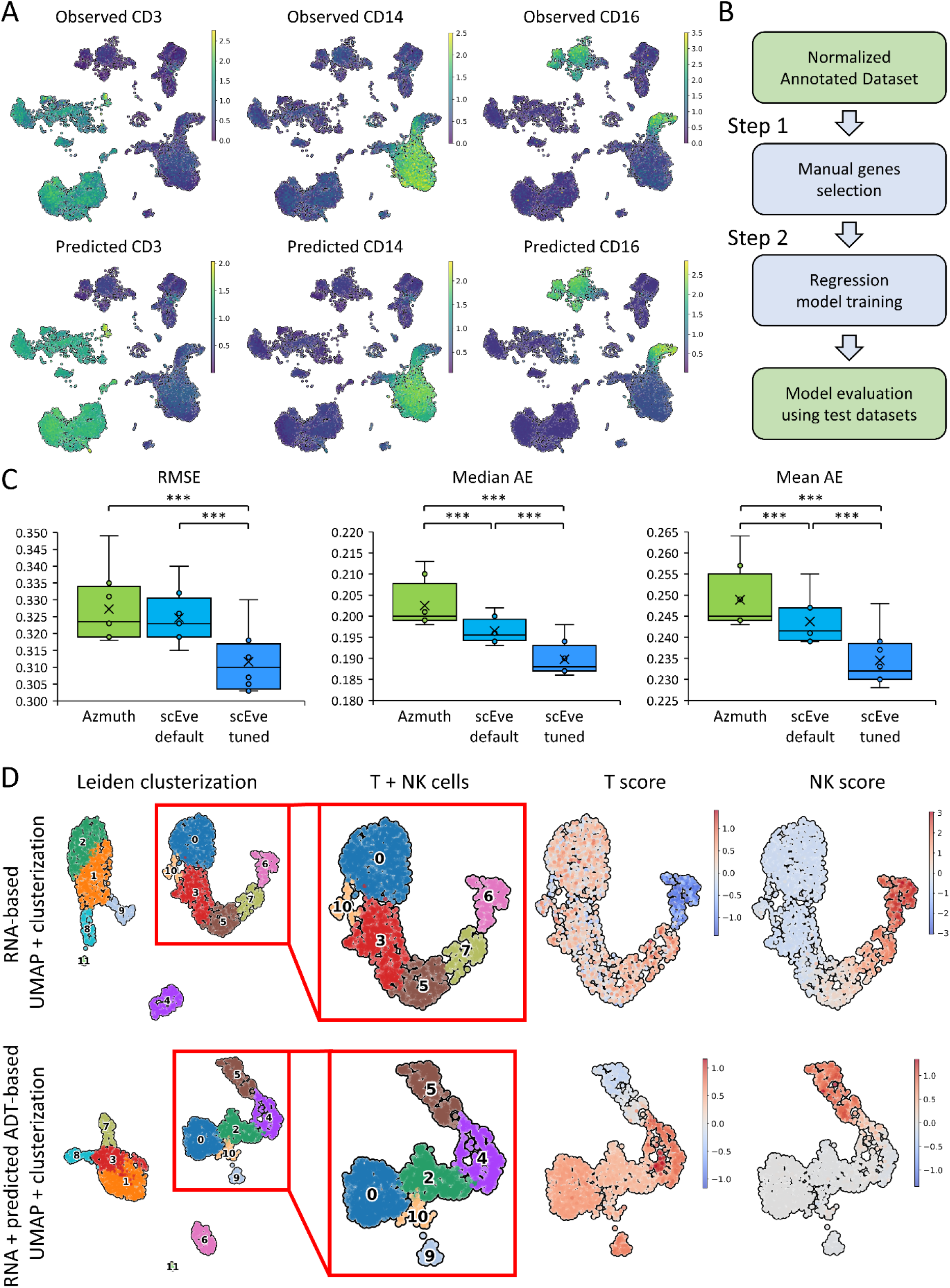
scEve model training and comparison with Azimuth cell surface data transfer. A. Observed and predicted cell surface proteins in the same cells. B. Pipeline of scEve model training and evaluation. C. Box plots comparison of Azimuth, scEve default and scEve tuned model-based prediction of cell surface proteins. RMSE - Root Mean Squared Error, AE - Absolute Error. For error metrics (RMSE, Median AE, Mean AE), a lower value indicates better prediction. D. Clusterization and UMAP calculated using only gene expression data (top row of pictures) and multimodal data - gene expression + predicted surface proteins (bottom row of pictures). T score (*CD3E, CD3D, CD3G* genes – top image; adt_CD3-1_pred, adt_CD3-2_pred – bottom image), NK score (*GNLY, NKG7, FCER1G, KLRF1* genes – top image; adt_CD56-1_pred, adt_CD56-2_pred, adt_CD158b_pred, adt_CD158_pred, adt_CD335_pred predicted proteins – bottom image). One Way RM ANOVA with post-hoc Tukey test. Median (line), mean (cross), n = 8 (dots), *** p<0.001.

To test the performance of the scAdam model, we compared it with other AI-based methods of automatic cell type annotation — celltypist and scGPT^4,5^. The same 8 training datasets were used to train these models. The remaining datasets were also used as test data. scAdam outperformed celltypist in all classification metrics for the l2 and l3 levels of annotation. There was no significant difference in the celltype_l1 annotation (Suppl. Fig. 3). scGPT had comparable accuracy to the scAdam model, but lower values for other classification metrics (Fig. 1G-H). Additionally, scAdam, unlike celltypist and scGPT, was able to predict all cell types in any test dataset. In all test datasets, comparable models were unable to detect AXL+SIGLEC6+ dendritic cells (ASDC). Moreover, celltypist and scGPT gave unstable results in the prediction of some cell types, such as CD8+ and CD4+ T proliferating cells, CD8+ T central memory cells, NK CD56bright cells, etc. (Suppl. Fig. 4-5). Thus, scAdam not only outperformed celltypist and scGPT but also achieved more reproducible results in cell annotation.

**Figure 3.**
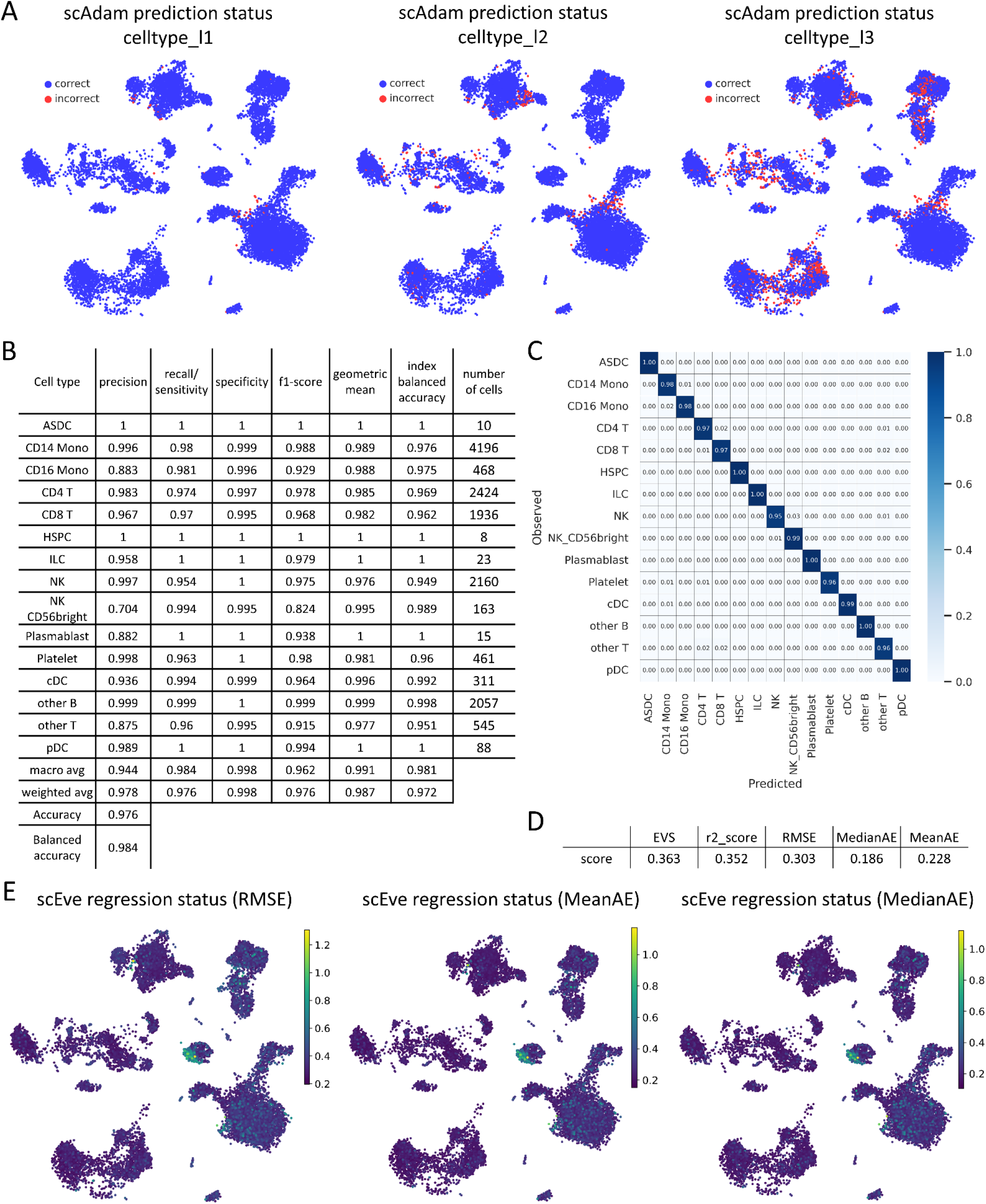
scNoah metrics for quality control of classification and regression tasks in scRNA-seq data. A. scAdam prediction status of cells (red – incorrectly predicted cell, blue – correctly predicted cells). B. Classification report of celltype_l2 annotation level (6 per cell type metrics + accuracy and balanced accuracy). C. Confusion matrix of prediction by cell type (celltype_l2). D. Report of cell surface protein prediction by scEve. RMSE – root mean squared error, MedianAE, MeanAE – median and mean absolute error. EVS – explained variance score, r2_score – coefficient of determination. E. scEve regression status. For error metrics (RMSE, Median AE, Mean AE), a lower value indicates better prediction.

### Cell surface protein abundance prediction using scEve

The second part of the scParadise set of tools was the scEve models. scEve made predictions of cell surface protein abundance based on the expression of selected features (Step 1 of the scAdam pipeline). As was seen in Figure 2A and Suppl. Fig. 6-7, the distribution of predicted cell surface proteins was comparable to that observed. To benchmark the performance of scEve predictions, we compared it with Azimuth^7^. To compare Azimuth and scEve, we used common regression metrics — root mean squared error (RMSE), mean and median absolute error (Mean AE, Median AE), coefficient of determination (r^2^ score), and explained variance score (EVS). All error metrics (RMSE, Mean AE, Median AE) worked on the principle that the smaller the value, the better the performance of the model. The default scEve outperformed Azimuth in Median AE, Mean AE, and r^2^ score. Hyperparameter tuning led to an increase in scEve performance in all regression metrics compared to the default model and Azimuth (Fig. 2C, Suppl. Fig. 7B).

Predicted cell surface proteins improved clusterization and annotation of transcriptionally similar cell types. One of the classic examples of the clusterization and annotation problem was the transcriptionally similar NK and CD8+ cytotoxic T cells. As shown in Figure 2D, the top row represented clusterization and cell embeddings calculated using only gene expression data. These clusters did not allow for a clear separation of T and NK cells based on marker gene expression (T, NK score - average expression of marker genes). Thus, cluster 7 included both NK (CD56+) and T (CD3+) cells (Suppl. Fig. 8 A,C). However, the use of predicted cell surface proteins improved clusterization and annotation of cell types. The bottom row represented cell embeddings and clusterization calculated using multimodal data (gene expression + predicted surface proteins) (Fig. 2D). Based on T and NK scores obtained using predicted surface proteins, it could be concluded that cluster 5 included NK cells, and cluster 4 included T cells. These data were supported by observed CD3 and CD56 proteins, as well as by visualization of gene-based and predicted protein-based T and NK scores (Suppl. Fig. 8). In addition, the prediction of surface proteins allowed us to distinguish CD8+ (cluster 9 on multimodal UMAP) and CD4+ (cluster 0 on multimodal UMAP) naïve T cells (Suppl. Fig. 9).

### Cell type classification and protein abundance prediction benchmarking using scNoah

To determine the performance of cell type annotation and modality prediction, we created the scNoah framework. First, scNoah allowed visualization of correct and incorrect annotations (Fig. 3A). Incorrect annotations made by automatic cell type annotation methods were mainly associated with transcriptionally similar cells, cell transition states, and rare cell types. An example of transcriptionally similar cells was B cell subtypes in the celltype_l3 annotation level of the PBMC dataset. Lambda and Kappa B cell subtypes differed only in the expression of immunoglobulin light chain genes — *IGLC1, IGLC2*, and *IGLC3* for B lambda cells and *IGKC* for B kappa cells. The remaining genes were expressed at approximately the same level. An example of annotation errors associated with cell transition states was CD14+ and CD16+ monocytes in the celltype_l2 (or celltype_l3) annotation level. CD14+ monocytes gave rise to CD16+ monocytes via an intermediate state called CD14+CD16+ intermediate monocytes^8,9^. Absolutely accurate annotation of such transition states was impossible, since changes in the cell transcriptome during differentiation occurred gradually. Small cell subpopulations often posed a challenge for AI-based methods due to the focus of models on obtaining high accuracy. However, we solved this problem within scAdam models. Second, scNoah allowed rapid application of key performance metrics for cell type annotation prediction, such as total accuracy, balanced accuracy, and per-cell type precision, sensitivity, F1-score, geometric mean, and index balanced accuracy of the geometric mean (Fig. 1B, Suppl. Fig. 10). In addition, scNoah allowed visualization of prediction performance using a confusion matrix (Fig. 2C, Suppl. Fig. 11 A-B).

To analyze the quality of cell surface protein abundance prediction, scNoah implemented the regression metrics RMSE, Mean AE, Median AE, r^2^ score, and EVS (Fig. 3D). In addition, all metrics could be visualized using cell embeddings, which allowed the detection of cells with low prediction quality. Using this method, we demonstrated that all metrics gave uniform prediction results among all cell types, except for a small proportion of platelets (Fig. 3E, Suppl. Fig. 11C).

Thus, scNoah was a set of tools for analyzing the quality of cell type and cell surface protein abundance prediction at the single-cell level.

## Discussion

We introduce scParadise, a publicly available Python package for tissue-specific cell type annotation (scAdam) and modality prediction (scEve). scAdam and scEve are fast deep learning-based methods. scAdam outperforms other AI-based annotation methods in annotating rare cell types representing less than 1% of the population, such as ASDC, ILC, HSPC, and T proliferating cells in PBMC datasets. This makes scAdam a unique tool for annotating rare cell types in scRNA-seq data. Another advantage of our model over other AI-based automatic cell type annotation methods is multitasking. This step facilitates the extraction of individual cell types from the dataset for more targeted investigation. The scEve models address another range of scRNA-seq problems, namely clusterization, visualization, and especially cell surface marker prediction. The prediction of cell surface protein markers allows for cell sorting to isolate a cell subpopulation of interest. The isolation of a cell subpopulation directs researchers toward a more targeted study using other methods such as ATAC-seq, microscopy, WB, and CRISPR-Cas9.

scParadise is also a tool for benchmarking cell type classification and modality prediction (scNoah). scNoah unifies the creation of automatic cell type annotation and modality prediction methods. For adequate comparison of methods, we recommend using several test datasets to check the reproducibility of predictions. Thus, we show that some automatic cell annotation methods, such as CellTypist and scGPT, can predict cell types inconsistently across different test datasets.

In the future, we plan to expand the number of models for cell type annotation and modality prediction. We also want to update our models through hyperparameter tuning and by increasing the number of cells in training datasets. In addition, we will update scParadise and add new methods and pipelines to speed up scRNA-seq data processing as well as to solve other problems related to this method.

## Methods

### Single-cell RNA-seq data analysis

#### PBMC CITE-seq dataset

The matrices of PBMC 3’ datasets were downloaded from GSE164378^6^. The dataset included 161,764 cells from 24 samples from 8 donors. We used corrected ready-made annotation of cells to increase the number of cells in small cell subsets. Also, we removed Erythrocytes and Doublets clusters from the dataset. We used scanpy and muon packages for downstream analysis. Gene matrix data was normalized using a shifted logarithm algorithm. We used centered log ratio (CLR) transformation (muon.prot.pp.clr) to normalize counts of cell surface protein labels. As the isotypes do not contain any biological information, we removed them from data. We used all protein labels and 2000 highly variable genes (scanpy.pp.highly_variable_genes(adata, n_top_genes = 2000, layer = ‘counts’, batch_key = ‘sample’, flavor = ‘seurat_v3’)) for separate gene and protein PCA analysis. We used multi-omics factor analysis (MOFA+), total variational inference (totalVI), weighted nearest neighbors (WNN) and PCA concatenation methods for multimodal data integration^6,10,11^. MOFA+, totalVI and WNN were used with default parameters. For PCA concatenation and WNN we used 20 principal components from each modality (RNA and protein) after batch correction using scANVI^12^. Batch correction of MOFA+ data was performed using harmony^13^. We used batch correction methods with default parameters and 2000 highly variable genes. Best method of multimodal data integration was defined by the scib package (WNN integration)^14^ (Suppl. Fig. 12, Suppl. table 1). All stages of dataset processing are available in the script PBMC_3p_dataset in scParadise repository (scripts_article folder).

#### Lung and Retina datasets

We downloaded processed Lung v2 (HLCA) and Retina datasets from CELLxGENE scRNA-seq data collection. We used ready-made annotation of cells.

### Feature selection for model training

We manually filtered highly variable genes for training and tuning machine learning models. We removed non-marker genes for any cell type (Suppl. Fig. 1A). We also added genes that were markers for cell types but were not included in the group of highly variable genes (Suppl. Fig. 1B). We used the sc.tl.rank_genes_groups function from the scanpy package and the FindAllMarkers function from Seurat with the Wilcoxon rank sum test to analyze cell type marker genes^15,16^. We mixed the resulting list of markers randomly to achieve an independent influence of genes on model training.

### Data split

We split the PBMC integrated dataset into training and 8 test datasets. For model training, we used 8 datasets. The test datasets included 2 samples from 2 donors. The training dataset was pruned based on the selected genes. The training dataset for the scAdam model was balanced for the average cell count across all cell types using the most detailed annotation level. An undersampling of major cell types was performed using random cell selection. An oversampling of rare cell types was performed using the RandomOverSampler function from the imblearn package with shrinkage=1. The training dataset for the scAdam and scEve models was split into a train dataset (90% of the training dataset) and a validation dataset (10% of the training dataset). The validation dataset was used for performance evaluation, preventing overfitting by early stopping, and also for hyperparameter selection.

### Machine learning model training for multi-task multi-class cell type annotation (scAdam)

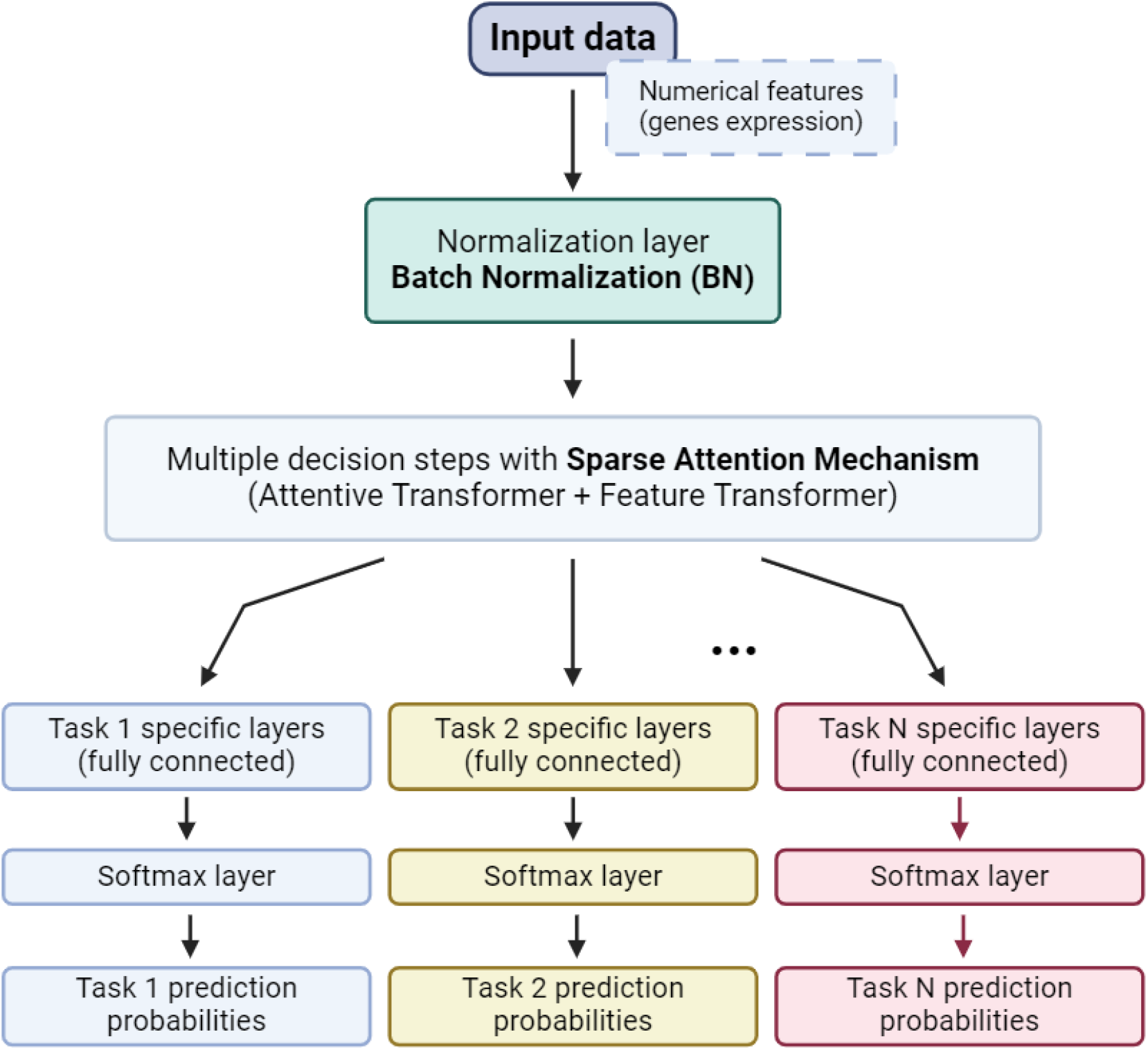

Scheme representing key points of scAdam structure.

Explanation of the Diagram

1. **Input Data (X)**: The raw input data containing both categorical and numerical features.
2. **Categorical Features**: Categorical features are processed through an embedding layer to convert them into dense vectors.
3. **Numerical Features**: Numerical features are normalized to standardize their scale.
4. **Shared Feature Transformer**: This component is shared across all tasks. It includes multiple decision steps, each using sparse attention to select relevant features dynamically.
5. **Task-Specific Branches**: After passing through the shared transformer, the data is split into multiple branches, each corresponding to a specific task. These branches consist of task-specific layers.
6. **Task-Specific Layers**: Each branch has its own fully connected layers to fine-tune the shared representations for the specific task.
7. **Output Heads**: Each task-specific branch ends with a softmax layer that outputs the class probabilities for the corresponding task.
8. **Predicted Probabilities**: The final outputs are the predicted class probabilities for each task.

In our study, we utilized TabNet, a neural network architecture developed by Google^17^. To address a multi-level annotation using tabular data we used a Multitask Learning version of TabNet - TabNetMultiTaskClassifier from PyTorch TabNet package. Unlike traditional neural networks, TabNet employs a sequential attention mechanism that allows the model to dynamically focus on different features at each decision step. This is crucial for multi-level prediction, as it enables the model to consider various aspects of the data at different stages of the decision-making process, leading to more nuanced and accurate predictions.

Additionally, TabNet’s architecture includes multiple decision steps, which allows the model to iteratively refine its focus and understanding of the features. This step-wise approach ensures that the model can progressively build a more detailed and accurate representation of the data, crucial for predicting multiple labels that may have complex interdependencies.

One of the most significant challenges in multi-level classification is understanding and interpreting the model’s decisions. TabNet addresses this challenge through its inherent interpretability. The model’s design allows for clear insights into which features are being utilized at each decision step. This high degree of interpretability allows us to check whether the model makes decisions that are adequate from the point of view of biological data.

The ability to scale and adapt to large data sets is critical in processing single-cell data. TabNet is designed to handle large volumes of data efficiently, making it suitable for big data applications. Its end-to-end learning capability means that the model can learn directly from raw input data to final output without extensive preprocessing or feature extraction. This adaptability simplifies the workflow with single-cell data.

### Multi-task regression model training for cell surface protein abundance prediction (scEve)

For solving the task of predicting the abundance of cell surface markers, we also used a modification of the TabNet structure - TabNetMultiTaskRegressor from the PyTorch TabNet package. For multitask regression, the TabNet architecture is modified to predict multiple continuous targets. The initial layers of the network are shared across all tasks. These layers capture common representations and patterns from the input data, leveraging shared learning. Following the shared layers, separate layers are dedicated to each regression task. These task-specific layers allow the model to fine-tune its predictions based on the unique characteristics of each target. The loss function in a multitask regressor is typically a weighted sum of the individual loss functions for each task. This ensures that the model optimally balances the performance across all tasks.

### Hyperparameters tuning of models

Hyperparameters tuning is crucial for optimizing the performance of machine learning models, including the TabNetMultiTaskClassifier and TabNetMultiTaskRegressor. For this task we used a tool such as Optuna. Optuna is an efficient and flexible hyperparameter optimization framework that can be used to automate this process.

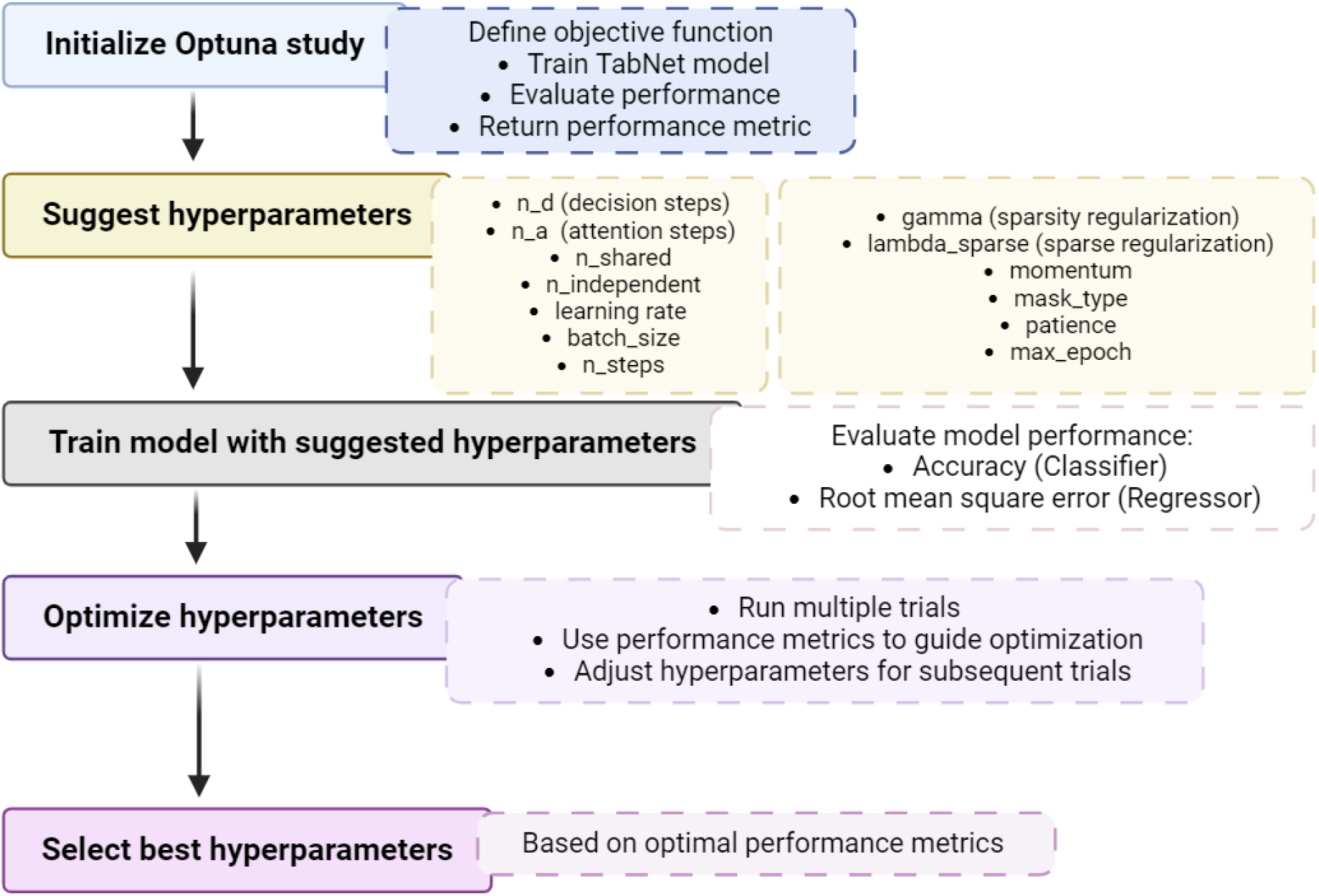

*Schematic Approach for Hyperparameters Tuning with Optuna*

1. Initialize Optuna Study Start by initializing an Optuna study, which will manage the hyperparameters optimization process.
2. Define Objective Function Define an objective function that trains and evaluates the TabNet model with different hyperparameters. This function is where the optimization will focus on finding the best set of hyperparameters.
3. Suggest Hyperparameters Within the objective function, use Optuna to suggest hyperparameters for each trial. These hyperparameters may include:

- n_d (number of decision steps)
- n_a (number of attention steps)
- learning_rate (the initial learning rate used for training)
- batch_size (number of examples per batch)
- virtual_batch_size (size of the mini batches used for “Ghost Batch Normalization”)
- n_steps (тumber of steps in the architecture)
- gamma (the coefficient for feature reusage in the masks)
- lambda_sparse (the extra sparsity loss coefficient)
- n_shared (number of shared Gated Linear Units at each step)
- cat_emb_dim (list of embeddings size for each categorical features)
- n_independent (number of independent Gated Linear Units layers at each step)
- momentum (momentum for batch normalization)
- mask_type (the masking function to use for selecting features)
- patience (number of consecutive epochs without improvement before performing early stopping)
- max_epochs (maximum number of epochs for training)
4. Train Model Train the TabNetMultiTaskClassifier or TabNetMultiTaskRegressor using the suggested hyperparameters.
5. Evaluate Performance Evaluate the model’s performance using a chosen metric, such as accuracy for classification or mean squared error for regression.
6. Optimize Hyperparameters Based on the performance metrics, Optuna will optimize the hyperparameters by running multiple trials. Each trial involves a different set of hyperparameters suggested by Optuna’s optimization algorithm.
7. Select Best Hyperparameters After completing all the trials, Optuna selects the best set of hyperparameters that resulted in the optimal performance.

### Benchmarking regression and cell type classification tasks in scRNA-seq (scNoah)

To create set of tools (scNoah) for analyzing cell type annotation performance, we used classic machine learning metrics (accuracy, balanced accuracy, precision, sensitivity, specificity, f1-score, geometric mean and index balanced accuracy of the geometric mean) from the scikit-learn and imblearn packages (scparadise.scnoah.report_classif_full)^18,19^. We also used a normalized confusion matrix to determine per cell type accuracy (scparadise.scnoah.conf_matrix). Additionally, we added visualization of correct and incorrect annotations on cell embeddings (scparadise.scnoah.pred_status). To assess the quality of cell surface abundance prediction, we used error metrics such as root mean squared error (RMSE), median and mean absolute error (MedianAE, MeanAE). Also, we used the explained variance regression score (EVS) and coefficient of determination (r2_score) to assess the quality of regression. These metrics can be applied both to the entire dataset to obtain average prediction values (scparadise.scnoah.report_reg) and to each cell separately (scparadise.scnoah.regres_status) for visualization on cell embeddings.

### Celltypist and scGPT

Celltypist and scGPT were used with default parameters. Models for the three annotation levels were trained using the same training dataset. The same testing datasets were used to analyze celltypist and scGPT performance. All steps of model training and performance testing are available in the “celltypist” and “scGPT” scripts in the scParadise repository (scripts_article folder).

## Data availability

Processed single cell RNA sequencing datasets were obtained from the CELLxGENE census (https://cellxgene.cziscience.com/datasets). The processed human lung scRNAseq dataset was retrieved from https://cellxgene.cziscience.com/collections/2f75d249-1bec-459b-bf2b-b86221097ced. The processed human retina scRNAseq dataset was accessed from https://cellxgene.cziscience.com/collections/4c6eaf5c-6d57-4c76-b1e9-60df8c655f1e. Matrices of CITE-seq PBMC 3’ scRNAseq were obtained from Gene Expression Omnibus (GSE164378)^6^. Matrices of multiome BMMC scRNAseq datasets were obtained from a single cell data integration challenge at the NeurIPS conference 2021 (https://openreview.net/forum?id=gN35BGa1Rt).

## Code availability

Code used for data analysis can be found on GitHub in scParadise repository: https://github.com/Chechekhins/scParadise/tree/main/scripts_article. You can find scripts for using scParadise in the same repository on GitHub: https://github.com/Chechekhins/scParadise/tree/main.

## Acknowledgements

We express our sincere gratitude to Anastasiya Y. Efimenko, who leads the Tissue Repair and Regeneration Laboratory at the Center for Regenerative Medicine, Medical Research and Education Institute, Lomonosov Moscow State University, as well as Pyotr A. Tyurin-Kuzmin and Konstantin Y. Kulebyakin from the Department of Biochemistry and Regenerative Biomedicine at the same institution for providing essential computing power that facilitated our research efforts.

