## Supplementary figures for "scParadise: Tunable highly accurate multi-task cell type annotation and surface protein abundance prediction"

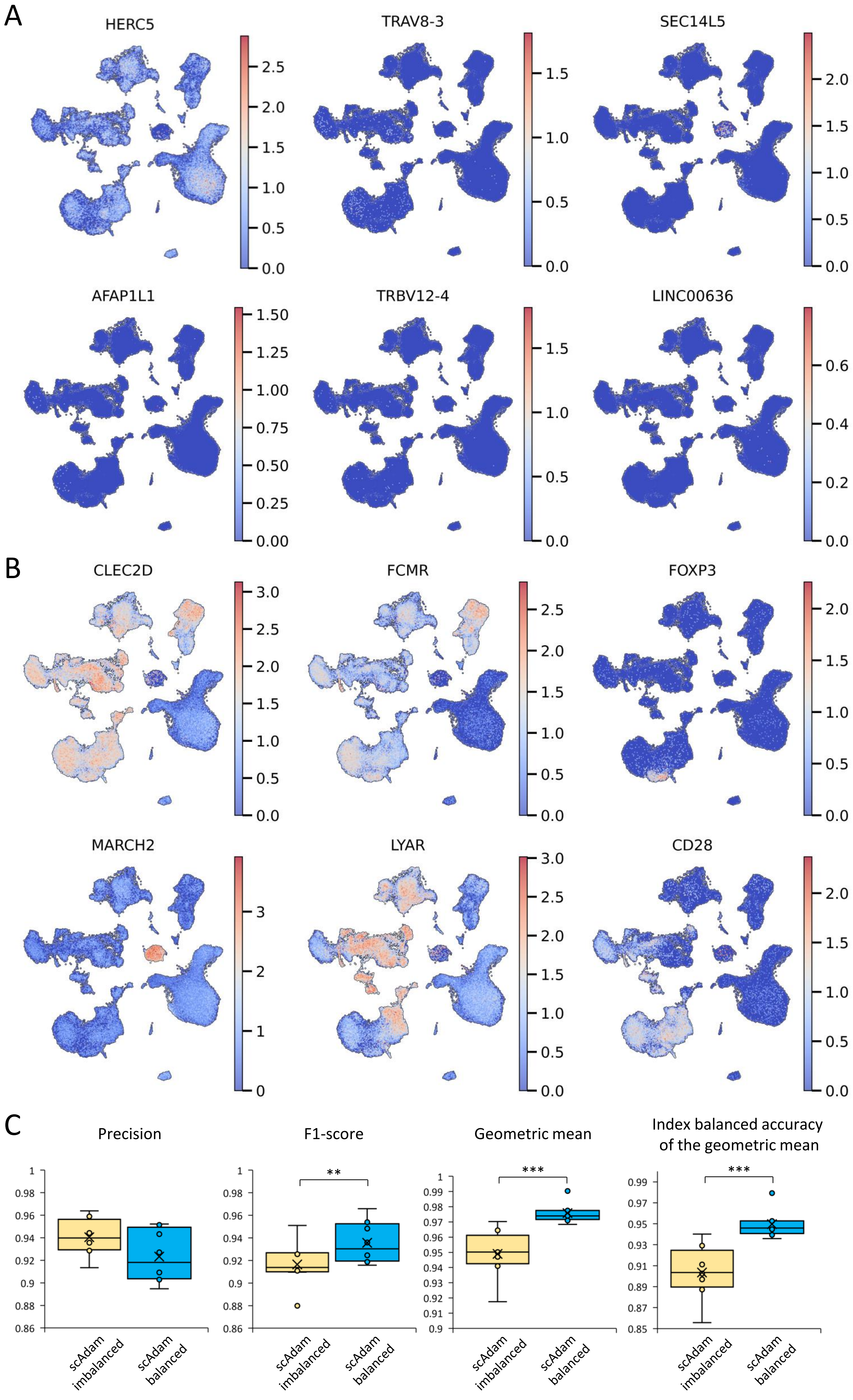


Supplementary Figure 1. Genes selection for scAdam and scEve models training. A. Examples of excluded highly variable genes. B. Examples of added marker genes not from list of highly variable. C. Comparison of scAdam imbalanced and balanced models prediction results.

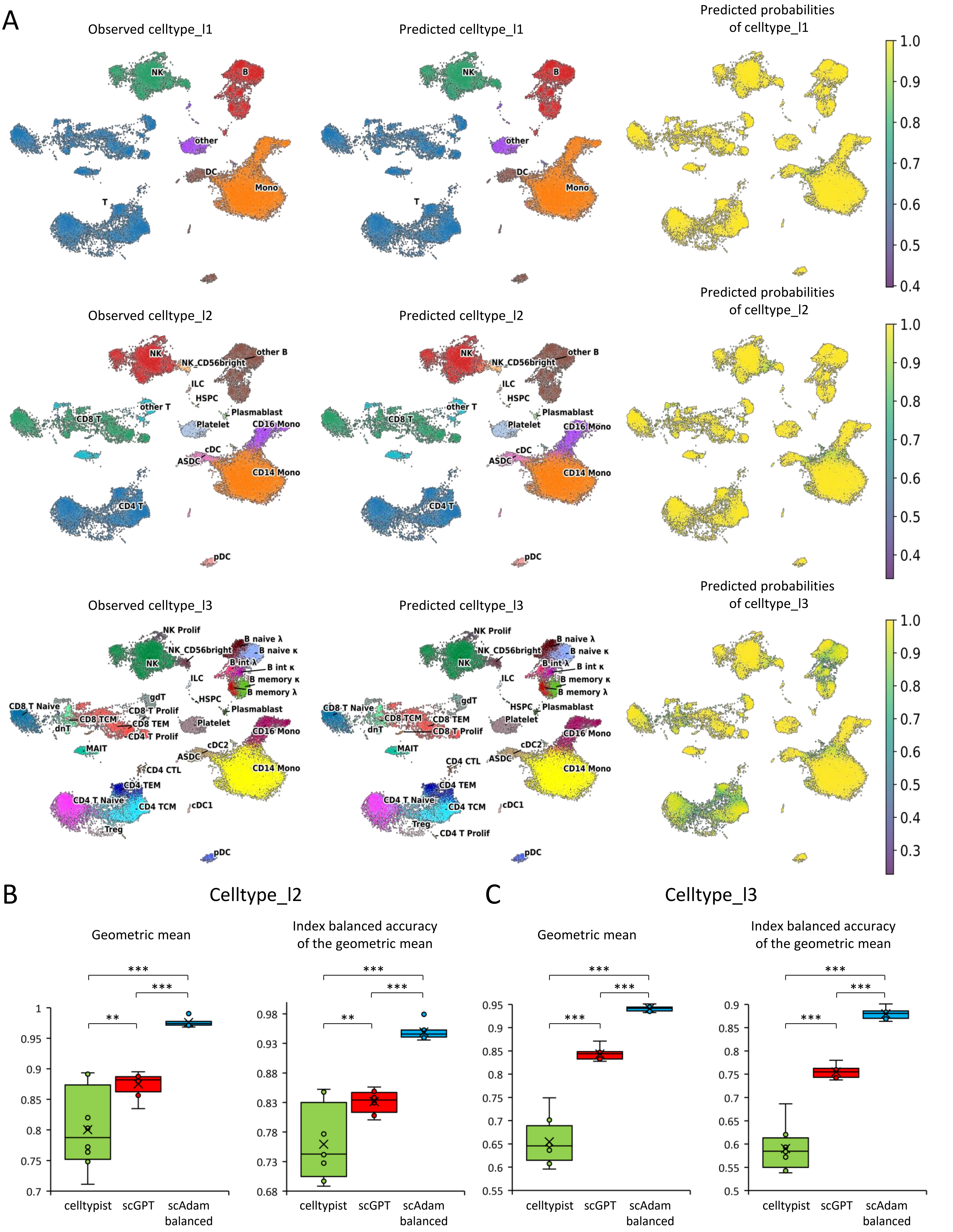


Supplementary Figure 2. scAdam prediction metrics. A. Results of scAdam prediction. The left column represents observed manually annotated cell types. The central column represents predicted cell types using scAdam model. The right column shows the probability that the cell belongs to the predicted type (higher value means higher probability of accurate annotation). B-C. Geometric mean and Index balanced accuracy of the geometric mean metrics of celltype_l2 (B) and celltype_l3 prediction (C).

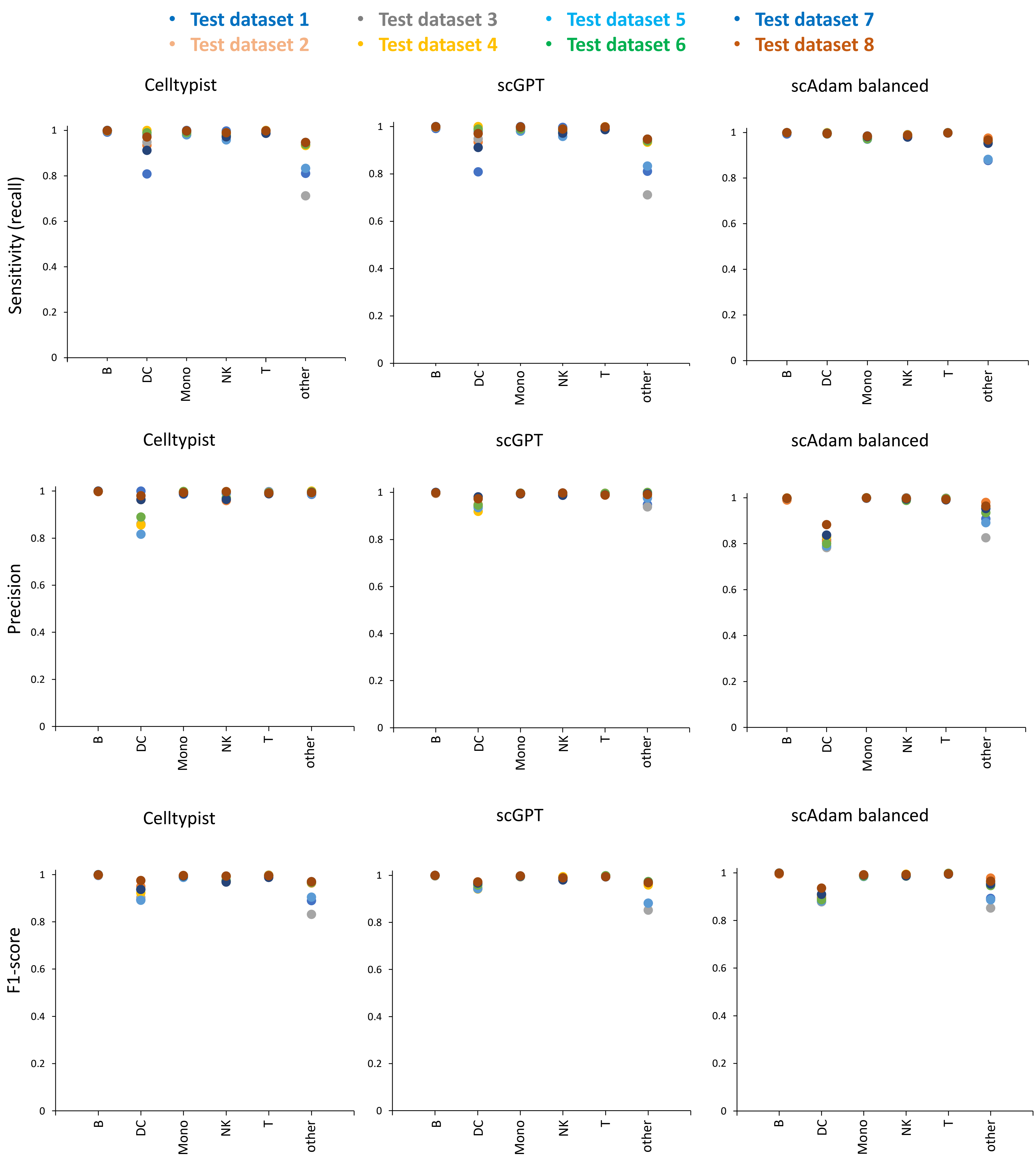


Supplementary Figure 3. Per cell type classification metrics of celltype_l1 annotation level.

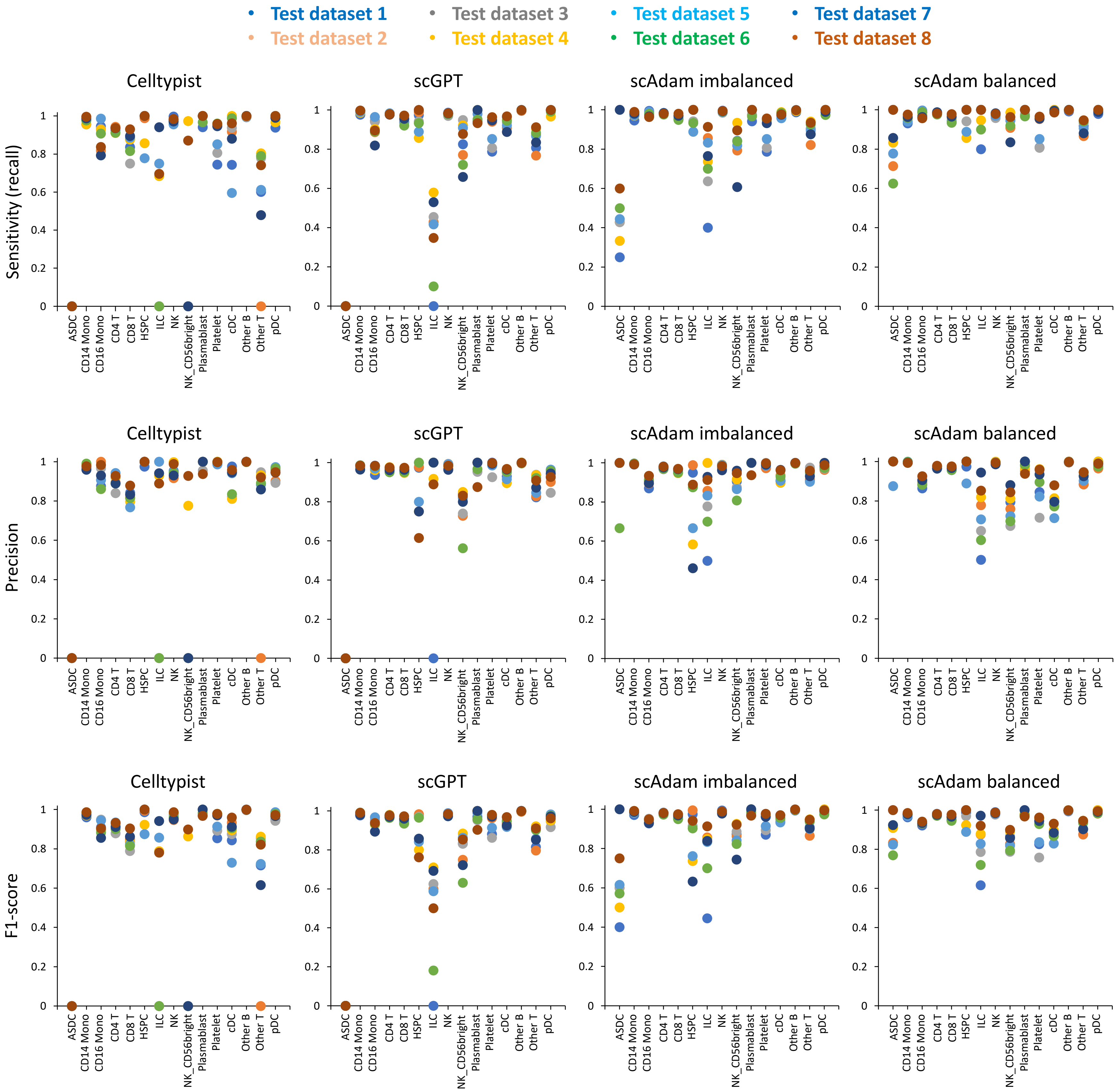


Supplementary Figure 4. Per cell type classification metrics of celltype_l2 annotation level.

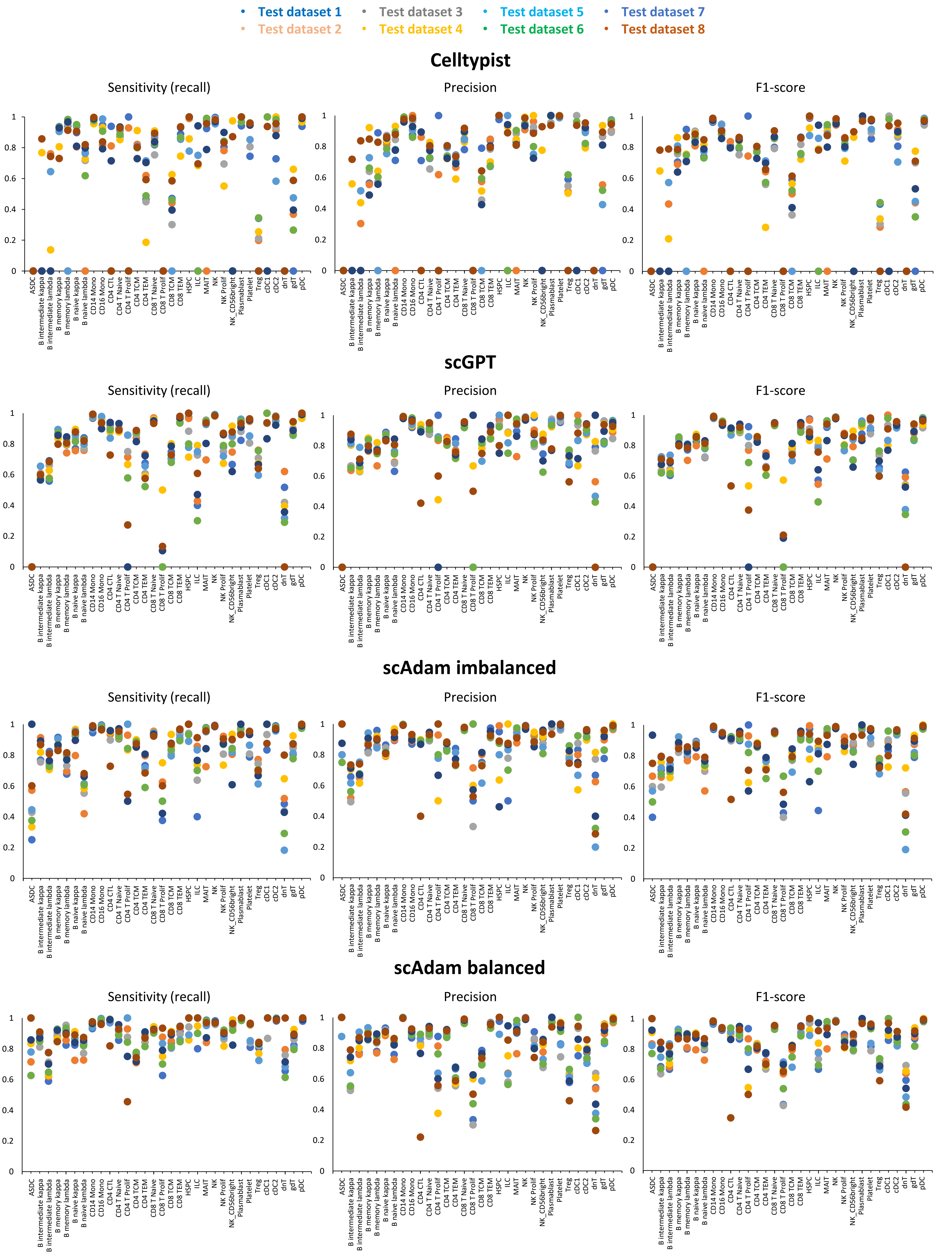


Supplementary Figure 5. Per cell type classification metrics of celltype_l3 annotation level.

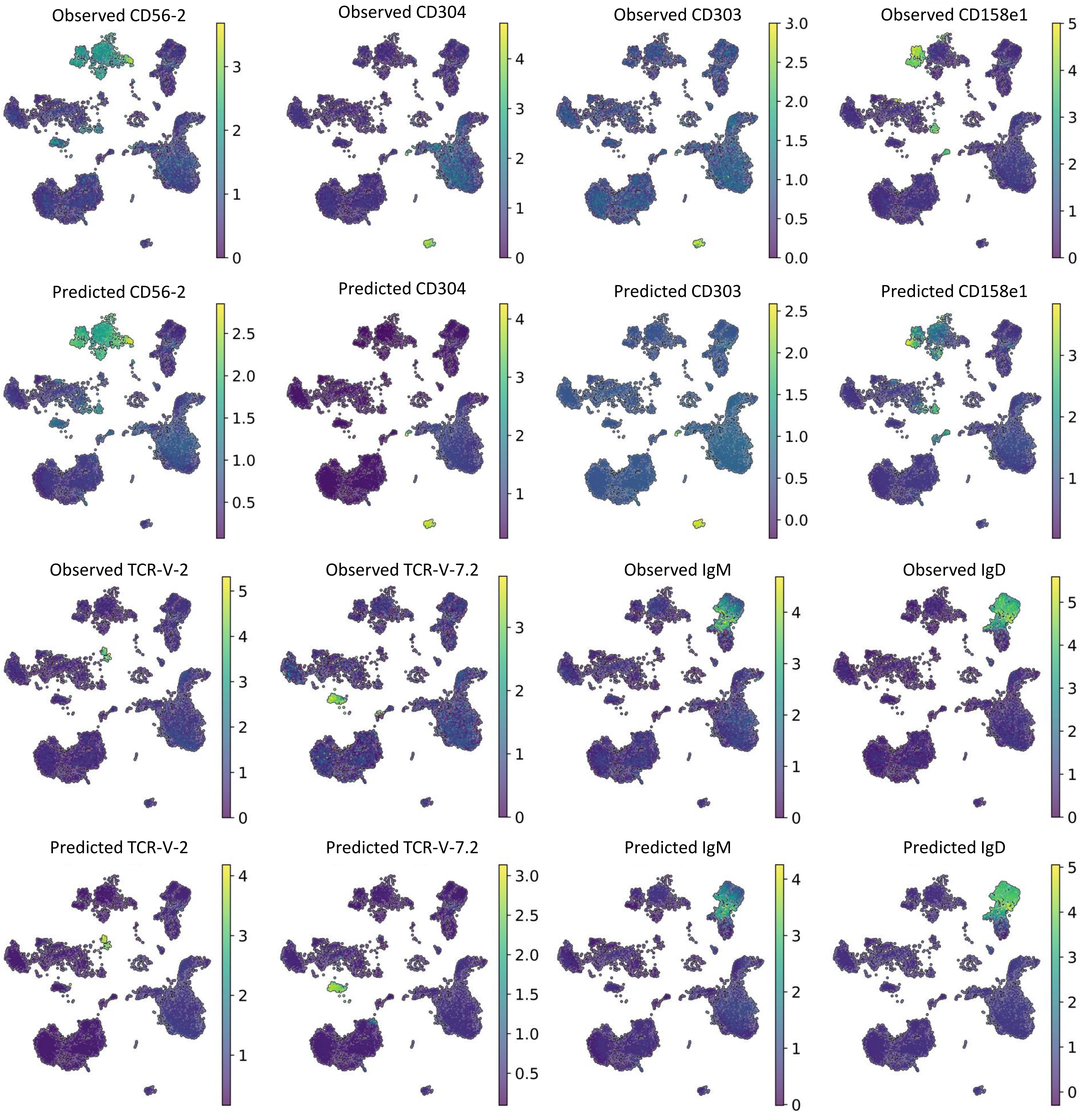


Supplementary Figure 6. Prediction of surface protein abundance using scEve models. Examples of predicted and observed surface proteins (antibody-1, antibody-2 – two separate antibodies with one target).

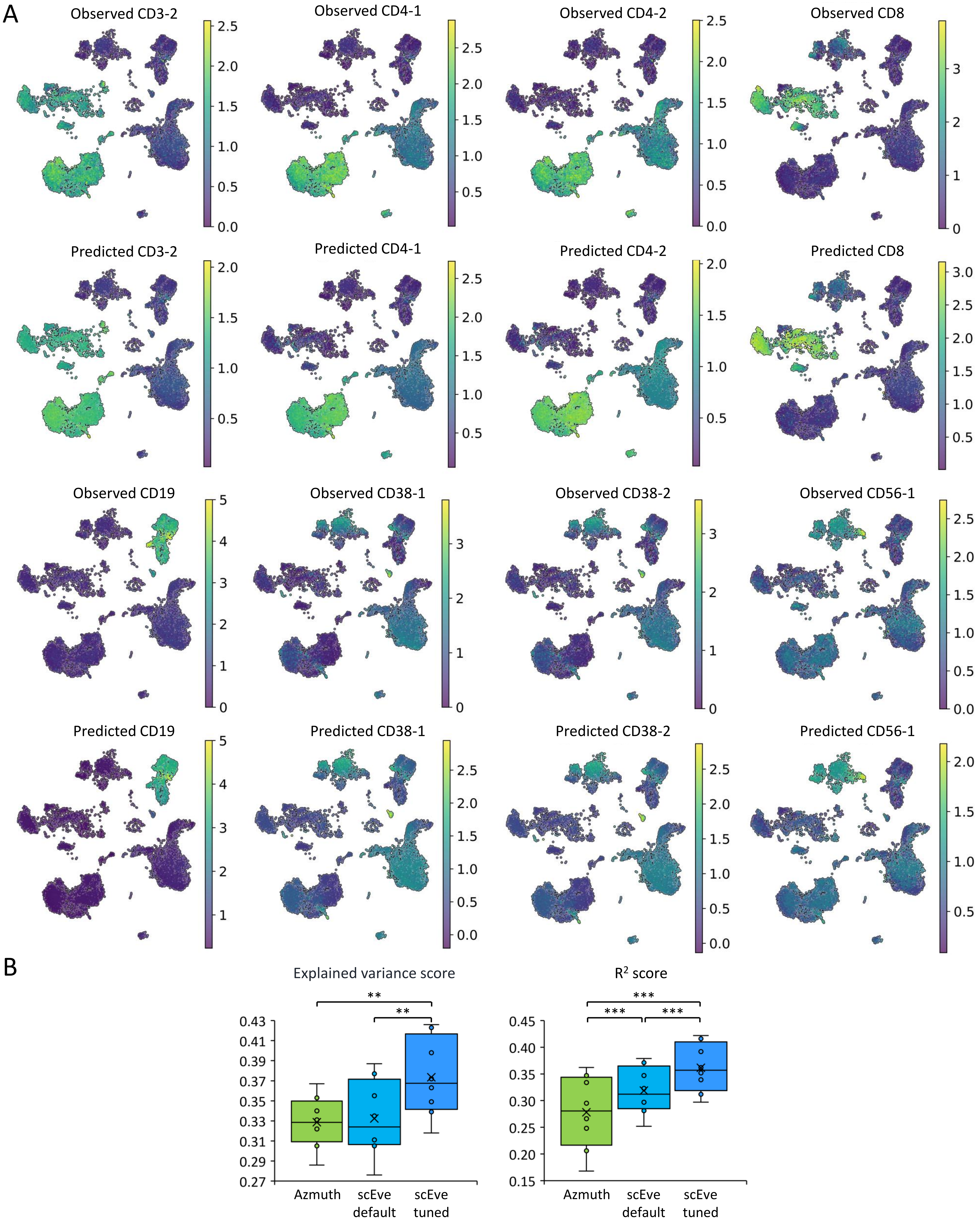


Supplementary Figure 7. Prediction of surface protein abundance using scEve models. A. Examples of predicted and observed surface proteins (antibody-1, antibody-2 – two separate antibodies with one target). B. Comparison of scEve default and tuned models with Azimuth results. One Way RM ANOVA with post-hoc Tukey test. Median (line), mean (cross), n = 8 (dots) , ** p<0.01, *** p<0.001.

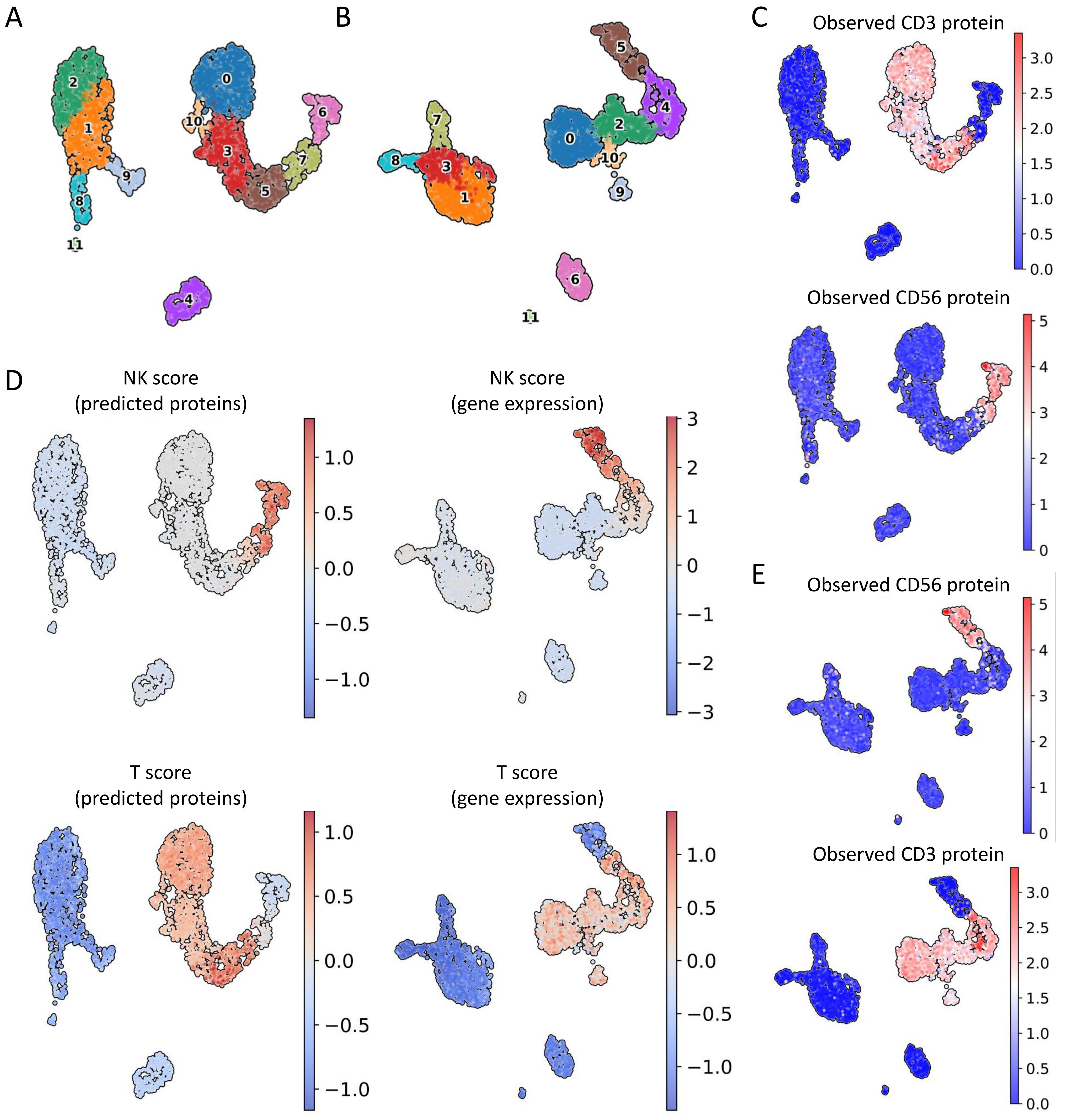


Supplementary Figure 8. NK and T scores on different UMAP plots. A. Clusterization and UMAP based on gene expression data. B. Clusterization and UMAP based on gene expression and predicted surface protein data. C. Observed CD3 and CD56 surface proteins on only gene expression-based cell embeddings (UMAP). D. T and NK scores based on gene expression data (NK score – *GNLY*, *NKG7*, *FCER1G*, *KLRF1*; T score – *CD3E*, *CD3G*, *CD3D*) and based on a predicted surface proteins (NK score – two subtypes of CD56 (CD56-1, CD56-2), CD158b, CD158, CD335; T score – two subtypes of CD3 (CD3-1, CD3-2)). E. Observed CD3 and CD56 surface proteins on multimodal (gene expression + predicted surface protein) data-based UMAP.

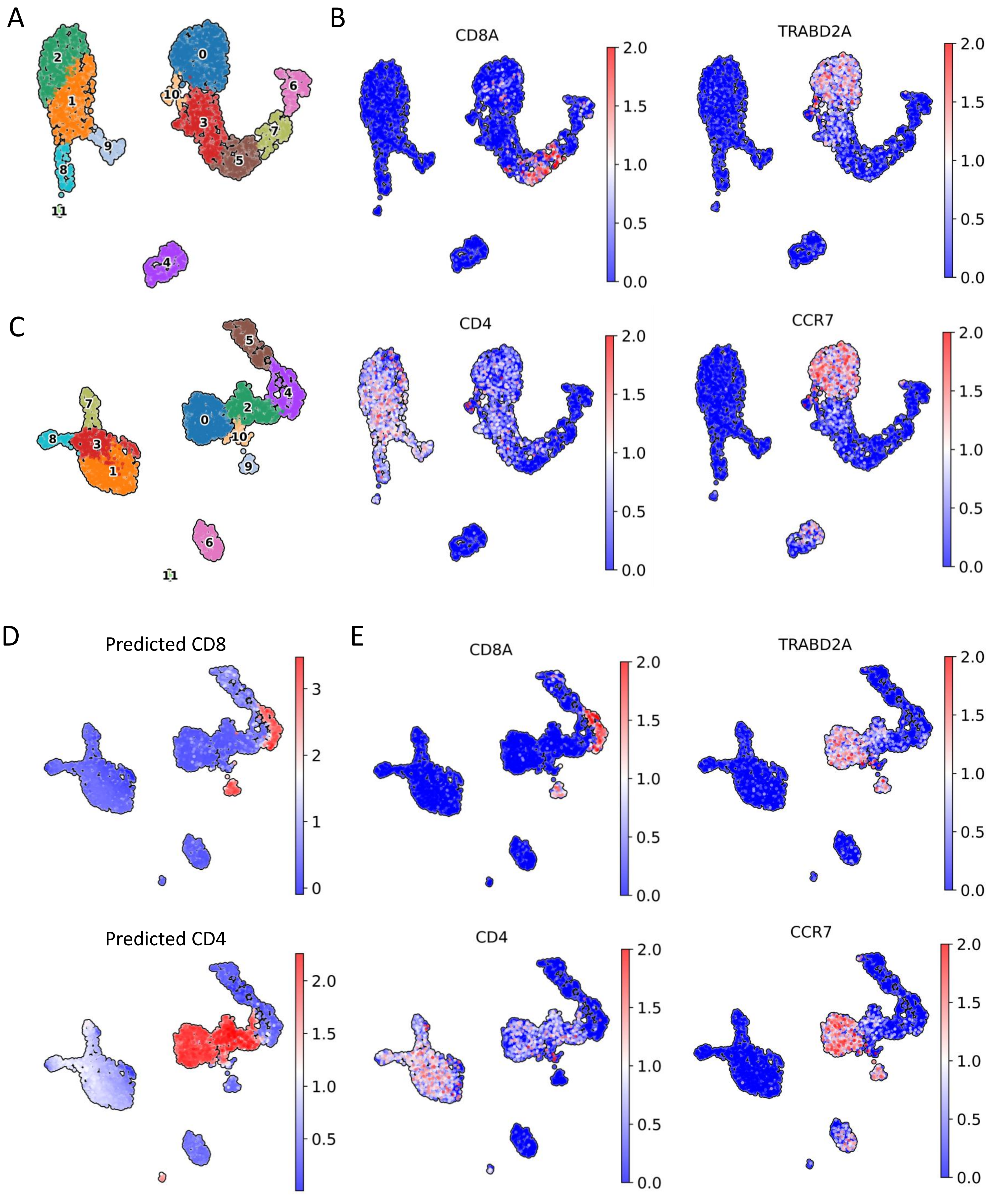


Supplementary Figure 9. Predicted surface proteins improves naïve T cells subtypes determination. A. Clusterization and UMAP based on gene expression data. B. Markers of naïve T cells (*CCR7*, *TRABD2A*) subtypes on UMAP calculated using gene expression data. C. Clusterization and UMAP based on gene expression and predicted surface protein data. D. Predicted CD8 and CD4 surface proteins. E. Markers of naïve T cells subtypes on UMAP calculated using gene expression and predicted surface protein data.

A

| Cell type | precision | recall/ sensitivity | specificity | f1-score | geometric mean | index balanced accuracy | number of cells |
|---|---|---|---|---|---|---|---|
| B | 0.998 | 0.999 | 1 | 0.999 | 0.999 | 0.999 | 2072 |
| DC | 0.951 | 0.993 | 0.999 | 0.971 | 0.996 | 0.991 | 409 |
| Mono | 0.998 | 0.995 | 0.999 | 0.997 | 0.997 | 0.994 | 4664 |
| NK | 0.997 | 0.986 | 0.999 | 0.991 | 0.993 | 0.984 | 2323 |
| T | 0.992 | 0.999 | 0.996 | 0.995 | 0.997 | 0.995 | 4905 |
| other | 0.996 | 0.967 | 1 | 0.981 | 0.984 | 0.964 | 492 |
| macro avg | 0.989 | 0.99 | 0.999 | 0.989 | 0.994 | 0.988 | |
| weighted avg | 0.994 | 0.994 | 0.998 | 0.994 | 0.996 | 0.992 | |
| Accuracy | 0.994 | | | | | | |
| Balanced accuracy | 0.99 | | | | | | |

B

| Cell type | precision | recall/ sensitivity | specificity | f1-score | geometric mean | index balanced accuracy | number of cells |
|---|---|---|---|---|---|---|---|
| ASDC | 1 | 1 | 1 | 1 | 1 | 1 | 10 |
| B int κ | 0.814 | 0.903 | 0.998 | 0.856 | 0.949 | 0.892 | 175 |
| B int λ | 0.828 | 0.877 | 0.998 | 0.852 | 0.935 | 0.864 | 187 |
| B memory κ | 0.931 | 0.798 | 0.999 | 0.859 | 0.893 | 0.781 | 252 |
| B memory λ | 0.866 | 0.897 | 0.998 | 0.882 | 0.946 | 0.887 | 195 |
| B naive κ | 0.929 | 0.868 | 0.997 | 0.897 | 0.93 | 0.854 | 721 |
| B naive λ | 0.827 | 0.899 | 0.993 | 0.862 | 0.945 | 0.885 | 527 |
| CD14 Mono | 0.996 | 0.979 | 0.998 | 0.988 | 0.989 | 0.976 | 4196 |
| CD16 Mono | 0.881 | 0.981 | 0.996 | 0.928 | 0.988 | 0.975 | 468 |
| CD4 CTL | 0.3 | 0.818 | 0.999 | 0.439 | 0.904 | 0.802 | 11 |
| CD4 T Naive | 0.933 | 0.909 | 0.995 | 0.921 | 0.951 | 0.897 | 1019 |
| CD4 T Prolif | 0.667 | 0.364 | 1 | 0.471 | 0.603 | 0.34 | 11 |
| CD4 TCM | 0.868 | 0.709 | 0.992 | 0.781 | 0.839 | 0.684 | 1036 |
| CD4 TEM | 0.509 | 0.818 | 0.984 | 0.628 | 0.897 | 0.792 | 297 |
| CD8 T Naive | 0.984 | 0.924 | 0.999 | 0.953 | 0.961 | 0.916 | 800 |
| CD8 T Prolif | 0.341 | 1 | 0.998 | 0.508 | 0.999 | 0.998 | 15 |
| CD8 TCM | 0.697 | 0.931 | 0.995 | 0.797 | 0.962 | 0.92 | 188 |
| CD8 TEM | 0.957 | 0.908 | 0.997 | 0.932 | 0.951 | 0.897 | 933 |
| HSPC | 1 | 1 | 1 | 1 | 1 | 1 | 8 |
| ILC | 0.958 | 1 | 1 | 0.979 | 1 | 1 | 23 |
| MAIT | 0.884 | 0.962 | 0.999 | 0.922 | 0.98 | 0.957 | 159 |
| NK | 0.996 | 0.947 | 0.999 | 0.971 | 0.973 | 0.941 | 2076 |
| NK Prolif | 0.811 | 0.869 | 0.999 | 0.839 | 0.932 | 0.857 | 84 |
| NK CD56bright | 0.73 | 0.994 | 0.996 | 0.842 | 0.995 | 0.99 | 163 |
| Plasmablast | 0.882 | 1 | 1 | 0.938 | 1 | 1 | 15 |
| Platelet | 0.998 | 0.959 | 1 | 0.978 | 0.979 | 0.955 | 461 |
| Treg | 0.554 | 0.82 | 0.998 | 0.661 | 0.905 | 0.804 | 50 |
| cDC1 | 0.941 | 1 | 1 | 0.97 | 1 | 1 | 16 |
| cDC2 | 0.93 | 0.99 | 0.998 | 0.959 | 0.994 | 0.987 | 295 |
| dnT | 0.294 | 1 | 0.999 | 0.455 | 1 | 0.999 | 5 |
| gdT | 0.874 | 0.932 | 0.996 | 0.902 | 0.964 | 0.922 | 381 |
| pDC | 0.989 | 1 | 1 | 0.994 | 1 | 1 | 88 |
| macro avg | 0.818 | 0.908 | 0.997 | 0.843 | 0.949 | 0.899 | |
| weighted avg | 0.933 | 0.922 | 0.997 | 0.925 | 0.958 | 0.913 | |
| Accuracy | 0.922 | | | | | | |
| Balanced accuracy | 0.908 | | | | | | |

Supplementary Figure 10. Classification report of celltype_l1 (A) and celltype_l3 (B) annotation level (6 per cell type metrics + accuracy and balanced accuracy)

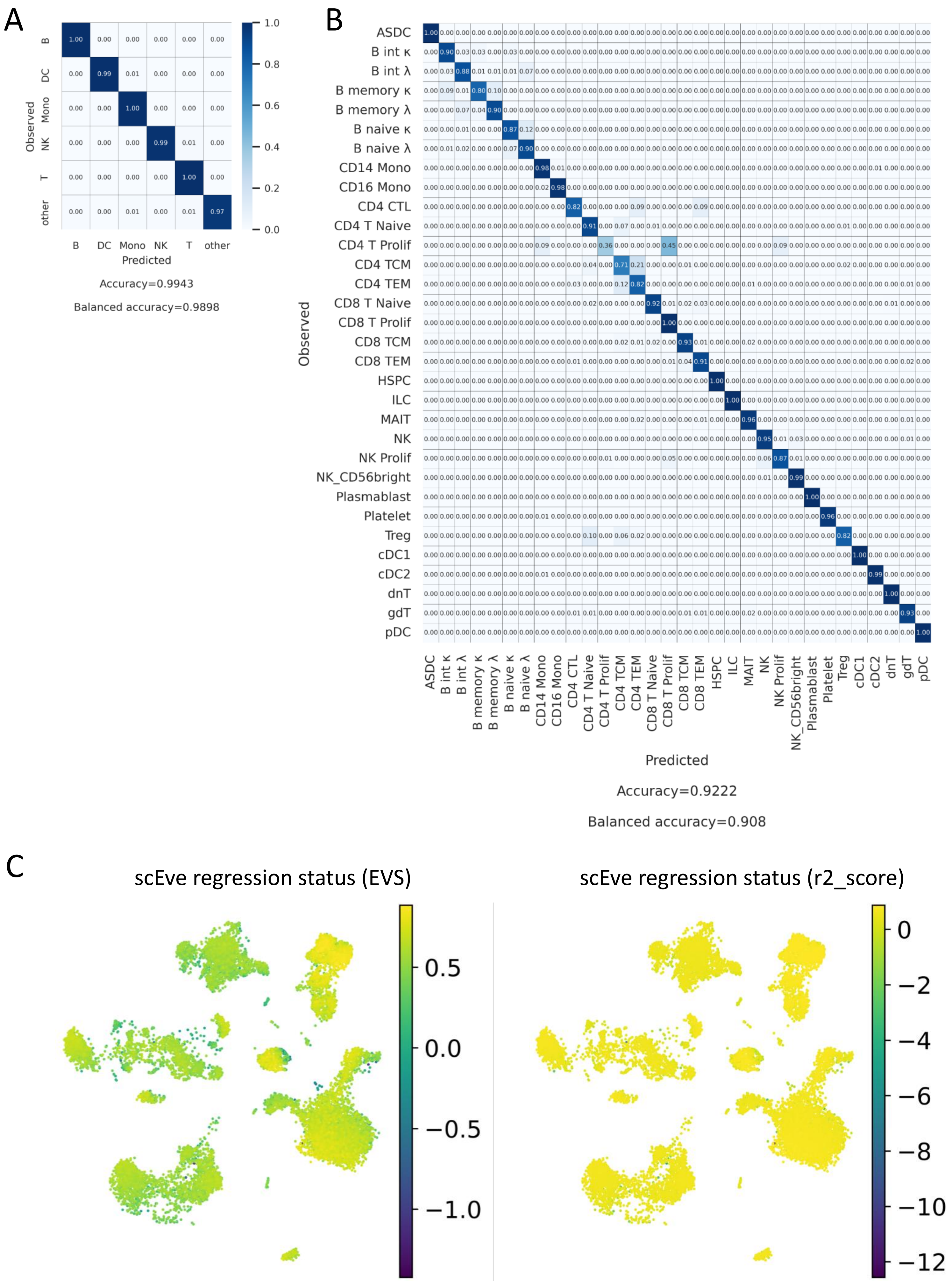


Supplementary Figure 11. scNoah confusion matrix for celltype_l1 (A) and celltype_l3 (B) annotation levels. C. Explained variance score (EVS) and coefficient of determination (r2_score) of scEve model prediction visualized on cell embeddings.

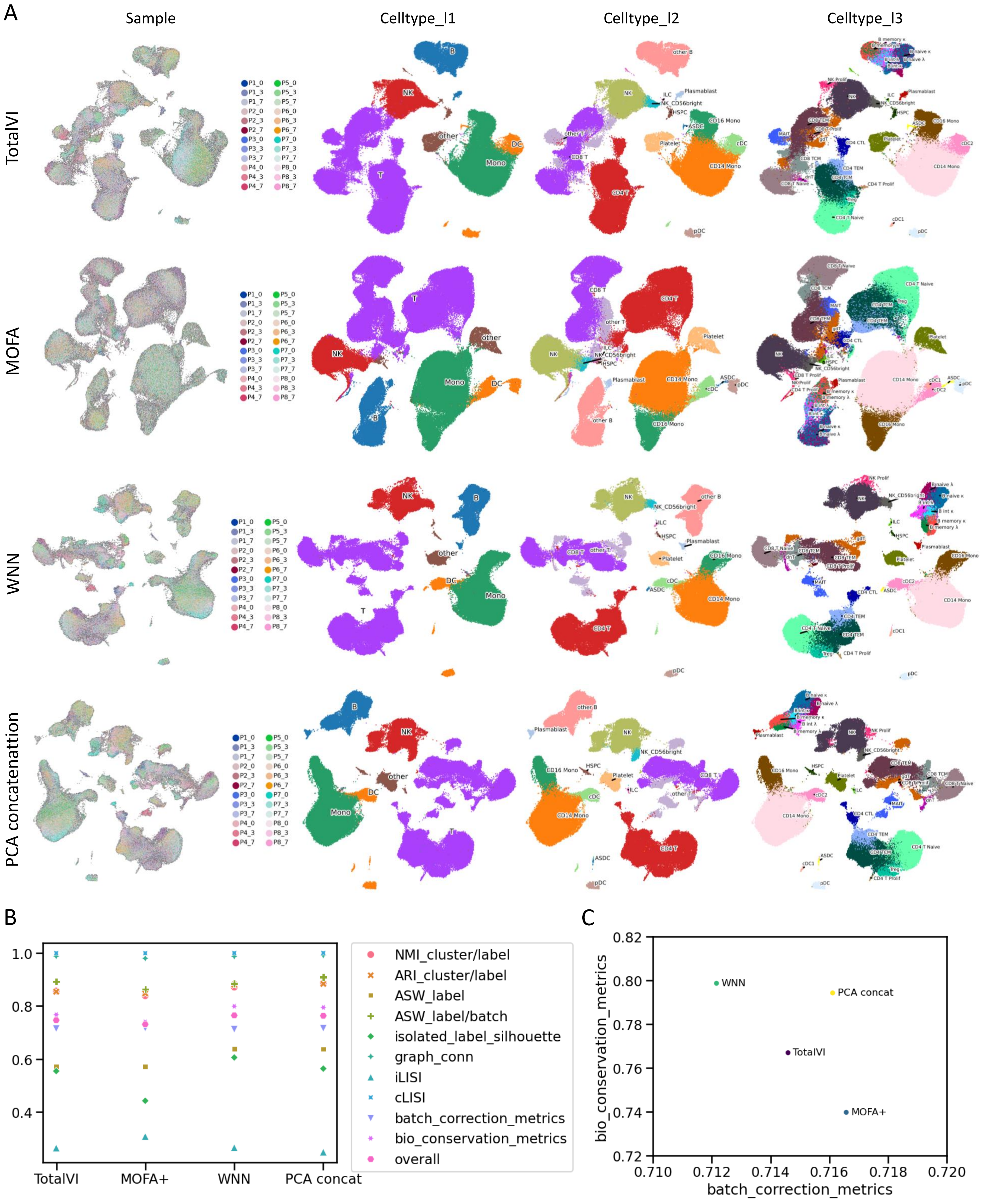


Supplementary Figure 12. PBMC CITE-seq data integration. A. Cell embeddings (UMAP) of different integration methods results. Cells colored by sample and 3 annotation levels - celltype_l1 (least detailed), celltype_l2, celltype_l3 (most detailed annotation level).
